# First-night effect reduces the beneficial effects of sleep on visual plasticity and modifies the underlying neurochemical processes

**DOI:** 10.1101/2024.01.21.576529

**Authors:** Masako Tamaki, Takashi Yamada, Tyler Barnes-Diana, Zhiyan Wang, Takeo Watanabe, Yuka Sasaki

**Affiliations:** Cognitive Somnology RIKEN Hakubi Research Team, RIKEN Cluster for Pioneering Research, Saitama, 351-0106, Japan; RIKEN Center for Brain Science, Saitama, 351-0106, Japan; Department of Cognitive, Linguistic, and Psychological Sciences, Brown University, Providence, RI, 02912, USA

## Abstract

Individuals experience difficulty falling asleep in a new environment, termed the first night effect (FNE). However, the impact of the FNE on sleep-induced brain plasticity remains unclear. Here, using a within-subject design, we found that the FNE significantly reduces visual plasticity during sleep in young adults. Sleep-onset latency (SOL), an indicator of the FNE, was significantly longer during the first sleep session than the second session, confirming the FNE. We assessed performance gains in visual perceptual learning after sleep and increases in the excitatory-to-inhibitory neurotransmitter (E/I) ratio in early visual areas during sleep using magnetic resonance spectroscopy and polysomnography. These parameters were significantly smaller in sleep with the FNE than in sleep without the FNE; however, these parameters were not correlated with SOL. These results suggest that while the neural mechanisms of the FNE and brain plasticity are independent, sleep disturbances temporarily block the neurochemical process fundamental for brain plasticity.

## Introduction

Understanding the roles of sleep is a central issue in neuroscience. The main roles of sleep include its effects on brain plasticity and learning. Performance on a task significantly improves after sleep ^1-7^, indicating the role of sleep in enhancing brain plasticity. Given that an estimated 50-70 million people suffer from sleep-related problems, including insomnia ^8,9^, it has become highly important to clarify how sleep disturbances influence brain plasticity.

However, this issue has not been thoroughly investigated. One of the obstacles hindering progress could be the presence of serious confounding factors when investigating the impacts of chronic sleep disturbances on brain plasticity. First, the effects of medications used for treatment, e.g., benzodiazepines and nonbenzodiazepines, could obscure the effects of sleep disturbances on learning ^10,11^. Second, chronic sleep disorders may be comorbid. Psychiatric symptoms, including depression and anxiety, are known to accompany chronic sleep disorders ^11^. It is difficult to dissociate sleep disorders and comorbid symptoms. In addition, medications for treating comorbid disorders are known to cause memory impairments ^12^. Thus, it has been a significant challenge to investigate the effects of sleep disturbance on brain plasticity without these confounding factors.

Here, we sought to investigate the effects of the first-night effect (FNE), a temporary sleep disturbance observed primarily during the first sleep session of experimental sessions ^13,14^, on brain plasticity. Individuals experiencing the FNE are known to suffer from delayed sleep onset, increased wake time after sleep onset and decreased time in deep sleep under the FNE ^15^. The FNE is not confined to nightly sleep but also manifests during daytime naps ^16^, and it is observed irrespective of individual anxiety levels among young and healthy individuals ^17^. Recent research has unveiled basic neural mechanisms associated with the FNE, indicating increased vigilance to monitor novel surroundings during sleep ^16,18,19^. However, how the FNE impacts on brain plasticity during sleep remains unclear.

This study aimed to fill this gap by investigating mechanisms underlying the effect of the FNE on plasticity. Because the FNE is mostly attenuated by the second sleep session ^15^, the FNE allows us to address whether and how disturbed sleep affects brain plasticity without having to account for the influence of confounding factors, such as medication and comorbidities. To address these questions, we compared disturbed sleep during the first session, which included the FNE (Day 1) with undisturbed sleep during the second session without the FNE (Day 2), in a within-subjects design among healthy participants.

We specifically examined *visual* plasticity, which has been extensively investigated in the human brain as an underlying mechanism of visual perceptual learning (VPL). VPL is defined as a long-term performance improvement on a visual task, including enhanced sensitivity to a visual feature ^20,21^. VPL is one of the learning types known to benefit from sleep ^4,7,22-25^. VPL improves significantly after sleep compared to before sleep without additional training – this phenomenon is termed offline performance gains ^1,7,26^. Thus, visual plasticity is measurable behaviorally by the amount of offline performance gains after sleep.

Moreover, visual plasticity can be measured neurochemically. Previous studies have shown that performance improvement in VPL is robustly represented by the regional excitatory-to-inhibitory neurotransmitter (E/I) ratio in early visual areas, and this ratio is measured by magnetic resonance spectroscopy (MRS) ^7,27-29^. *Increases* in the E/I ratio from baseline are significantly correlated with increased visual plasticity, shown as increased performance of VPL during wakefulness ^27,28^. A close association between enhanced visual plasticity and an increased E/I ratio was observed not only during wakefulness but also during sleep. A recent study ^7^ revealed that the E/I ratio during non-rapid eye movement (NREM) sleep was greater than the baseline E/I ratio during wakefulness. This finding indicates that the visual system became more plastic during NREM sleep than during wakefulness. Moreover, the increases in the E/I ratio during NREM sleep were robustly correlated with offline performance gains in VPL. These results suggest that the degree of visual plasticity associated with sleep could be measured by both how much offline performance gains in VPL occur with sleep and how much the E/I ratio increases during sleep.

Given our understanding of the effects of normal sleep on plasticity, the present study focused on investigating how the FNE impacts visual plasticity in two independent nap experiments using two different outcomes, i.e., offline performance gains and increases in the E/I ratio during sleep. We employed a state-of-the-art neuroimaging method of simultaneous MRS and polysomnography (PSG) measurements to evaluate changes in the E/I ratio during sleep ^30^. The results showed that the FNE disrupts offline performance gains and impedes neurochemical changes that are fundamental for offline performance gains. These results provide evidence that even a transient decrease in sleep quality among healthy adults may negatively impact brain plasticity during sleep.

## Results

### Experiment 1

In Experiment 1, a group of young, healthy participants (n=17) participated in two sessions (Day 1 and Day 2) in the laboratory (**Fig. 1A**, see details in the Methods section). To explore how the FNE impacts offline performance gains during sleep, we compared the offline performance gains over the nap session between Days 1 and 2 (**Fig. 1A**). We used the texture discrimination task (TDT; see details in the Methods section). The VPL of a TDT is specific to the visual location of the trained target. Namely, the VPL of a TDT with a target presented in one visual location (for instance, the left visual location) does not transfer to untrained visual locations (for instance, the right visual location). Thus, if two different visual locations are used between Days 1 and 2, then learning should occur independently between these two days, allowing us to measure offline performance gains for Days 1 and 2 independently within the same participants.

**Fig. 1.**
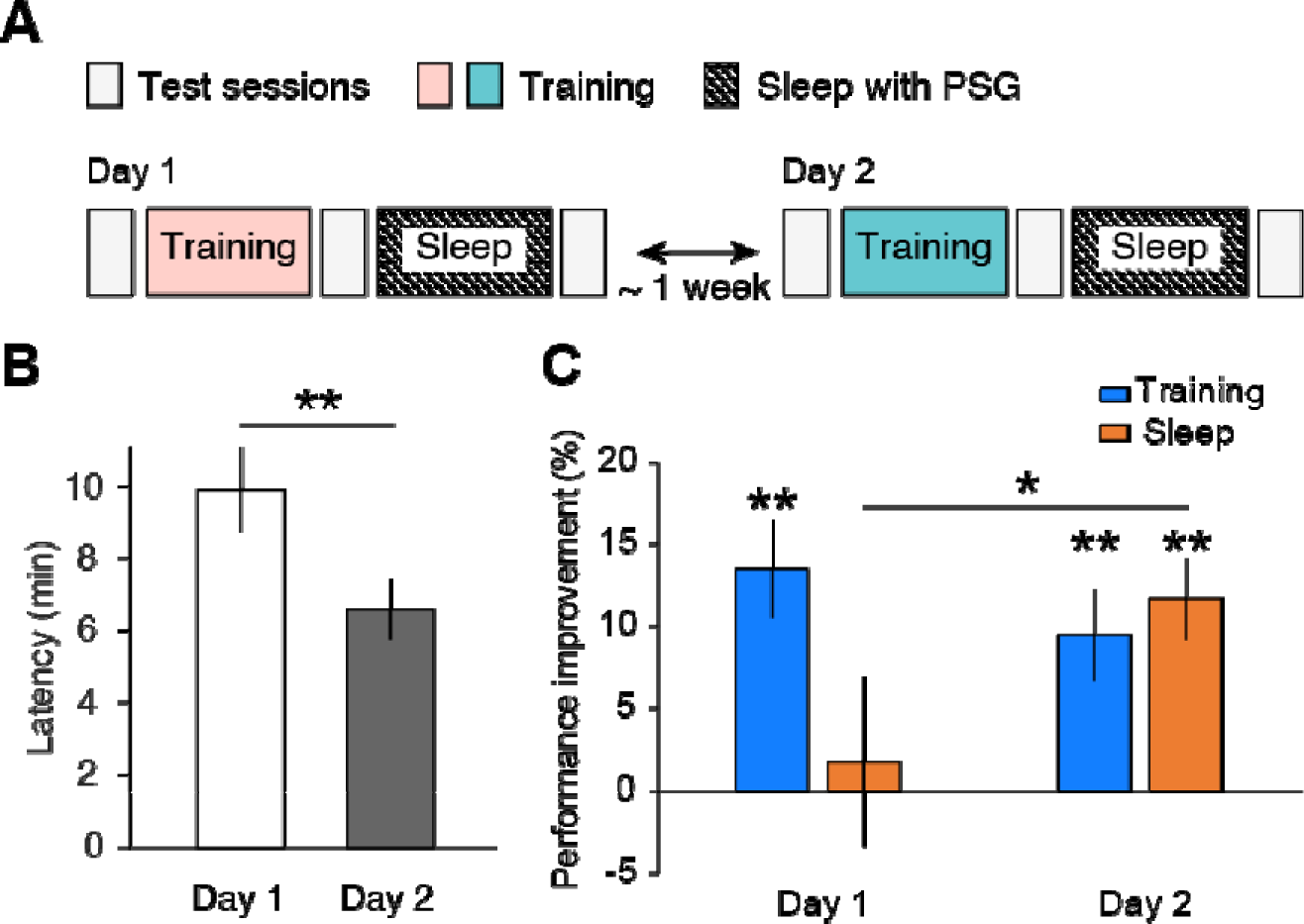
Experiment 1. **A.** Experimental design. The pink and teal boxes show different visual hemifields trained between Days 1 and 2. **B.** Sleep onset latency. **C.** Performance improvement after training (cyan) and after sleep (orange) on Days 1 (left bars) and 2 (right bars). One-sample t test or paired t test, **p<.01, *p<.05. N=17.

On Days 1 and 2, the participants were asked to perform training on the TDT, where the target appeared in a consistent visual hemifield (the left or right) before the nap session. The participants were asked to fixate at the center and to report the orientation of the target (horizontal or vertical) in the TDT. The trained visual hemifield (left or right) was randomly assigned to a subject on Day 1 and switched on Day 2. The trained visual hemifield on Day 2 was the untrained visual hemifield on Day 1. There were three test sessions: pretraining, posttraining (which occurred before sleep), and postsleep sessions (**Fig. 1A**). The behavioral measure was the threshold stimulus-to-ask onset asynchrony (SOA in ms), at which participants achieved 80% accuracy in the orientation task. Performance changes were calculated based on the threshold SOAs. See the Methods section for more details.

We tested whether the FNE occurred on Day 1 (**Table 1**) by using sleep onset latency (SOL, min) in accordance with previous studies ^16,19^. The nonparametric Wilcoxon signed-rank test was used due to the nonnormal distribution of the sleep variables (**Table 1**). The results showed that the SOL was significantly different between days (**Fig. 1B**, *Z*=3.115, n=17, *p*=0.002, 95% CI=[1.6, 5.6]). The SOL was significantly longer on Day 1 than on Day 2, providing evidence that the FNE occurred in Experiment 1.

**Table 1:** Sleep parameters for Experiment 1.

|  | Day 1 |  | Day 2 |  | Wilcoxon signed rank test |  |
| --- | --- | --- | --- | --- | --- | --- |
|  |  |  |  |  | Z | P |
| SOL (min) | 9.9 | ± 1.18 | 6.6 | ± 0.85 | 3.12 | 0.002 |
| WASO (min) | 10.0 | ± 2.62 | 4.7 | ± 1.93 | 2.28 | 0.023 |
| Stage W (%) | 19.7 | ± 3.91 | 10.1 | ± 2.55 | 2.77 | 0.006 |
| Stage N1 (%) | 13.3 | ± 1.44 | 14.3 | ± 2.25 | -0.36 | 0.723 |
| Stage N2 (%) | 34.5 | ± 2.12 | 38.6 | ± 2.98 | -1.21 | 0.227 |
| Stage N3 (%) | 25.9 | ± 4.76 | 30.7 | ± 4.30 | -1.54 | 0.124 |
| Stage R (%) | 8.1 | ± 2.19 | 12.4 | ± 2.60 | -1.35 | 0.177 |
| Total time (min) | 81.9 | ± 1.47 | 77.7 | ± 3.54 | 0.57 | 0.569 |

SOL, sleep-onset latency. WASO, wake after sleep onset. SE, sleep efficiency. The duration of each sleep stage was measured from the first sleep cycle. The total time was defined as the sum of W, N1, N2, N3, and R stage durations within the first sleep cycle. Values are means ± SEs. Wilcoxon signed-rank test (n=17) was applied to test whether sleep parameters were significantly different between sleep sessions (days 1 vs. 2).

We next examined whether the expected offline performance gains in TDT performance were attenuated by the FNE. If the FNE impairs visual plasticity, the degree of offline performance gains during the sleep session may be significantly lower, leading to a significant difference between Days 1 and 2. On the other hand, if the FNE does not impact visual plasticity, there should be no significant difference in the degree of offline performance gains between Days 1 and 2. We calculated the relative performance changes over the training (the change in the threshold SOAs from pre-to posttraining, divided by the pretraining SOA) and offline performance gains (the change in the threshold SOAs from posttraining to postsleep, divided by the posttraining SOA) on each day.

Repeated-measures 2-way ANOVA with the factors Day (Day 1 and 2) and Session (training and sleep) was performed to evaluate changes in TDT performance. There was a significant interaction effect between the two factors (F(1,16)=4.67, p=0.046), while the main effects of Day (F(1,16)=0.67, p=0.425) and Session (F(1,16)=2.42, p=0.139) were not significant (**Fig. 1C**).

Next, to investigate the source of the interaction, we tested whether there was a significant difference in the performance changes due to training and offline performance gains between days. While the amount of performance change due to training was not significantly different between days (paired t test, t(16)=0.77, p=0.451, 95% CI=[-7.0, 15.0]), the quantity of offline performance gains was significantly different between days (paired t test, t(16)=-2.21, p=0.042, 95% CI=[-19.5, -0.4]; **Fig. 1C**). Moreover, we tested whether the performance changes due to training and the offline performance gains by sleep were significant each day. On both days, significant performance improvements occurred after training (Day 1, one-sample t test against zero, t(16)=4.51, p<0.001, 95% CI=[7.1, 19.8]; Day 2, one-sample t test against zero, t(16)=3.42, p=0.004, 95% CI=[3.6, 15.4]). On the other hand, significant offline performance gains occurred only on Day 2 (one-sample t test against zero, t(16)=4.68, p<0.001, 95% CI=[6.4, 17.0]) and not on Day 1 (one-sample t test against zero, t(16)=0.34, p=0.736, 95% CI=[-9.2 12.7]). These results are consistent with the hypothesis that the FNE disrupts visual plasticity during sleep.

We conducted control analyses to test whether the larger offline performance gains on Day 2 were due to interactions between the two independent training sessions. First, we tested whether unexpected learning transfer occurred from Day 1 to Day 2. If learning was transferred from Day 1 to Day 2, the initial performance level should be significantly better on Day 2 than on Day 1. However, the threshold SOA at the pretraining test session was not significantly different between Days 1 and 2 (paired t test, t(16)=1.40, p=0.182, 95% CI=[-7.2, 34.8]). Next, we tested whether learning on Day 1 anterogradely interfered with that on Day 2. If interference occurred on Day 2, the performance level at the posttraining test session would be worse on Day 2 than on Day 1, leading to larger offline performance gains on Day 2. However, a paired t test showed no significant difference between the threshold SOA at the posttraining test session between days (t(16)=0.67, p=0.513, 95% CI=[-9.2, 17.7]). These results are consistent with the above ANOVA results that the amounts of performance improvements due to training were not significantly different between Days 1 and 2, suggesting that learning on Days 1 and 2 was independent and that it is unlikely that transfer or interference between the two training sessions would result in a significant amount of offline performance gains on Day 2.

In addition, we tested whether any factors other than the FNE had an effect on the degree of offline performance gains on Day 1, as in Experiment 1. Sleepiness was measured by the psychomotor vigilance task (PVT) ^31^ and the Stanford Sleepiness Scale (SSS) ^32^ prior to each of the test sessions. Since both the PVT and SSS deviated from the normal distribution, including outliers, we used the Wilcoxon signed-rank test. We tested whether the lack of significant offline performance gains on Day 1 was caused by the increased sleepiness in the test sessions on Day 1. However, neither sleepiness test indicated that this was the case (see **Table 2** for the results of the statistical tests) as follows. First, the PVT data were not significantly different between Days 1 and 2 at any of the test sessions. Second, the SSS scores were not significantly different at the pretraining and postsleep test sessions between Days 1 and 2, but there were significant differences at the posttraining session, which was before the sleep session (**Table 2**). However, this difference in the SSS score at the posttraining session does not account for the smaller offline performance gains on Day 1. While the range of SSS scores at the posttraining session was significantly smaller on Day 2 than on Day 1, resulting in a significant difference according to the Wilcoxon signed-rank test result, the median SSS score was the same (scored as 2) on both days. The SSS scores are discrete values, and a score of 2 indicates that the subject felt highly functional, not drowsy ^32^. Thus, these data suggest that sleepiness was not significantly different between Days 1 and 2 and that sleepiness does not account for the small offline performance gains on Day 1.

**Table 2.** Sleepiness. Reaction times (RTs) on a psychomotor vigilance task (PVT) and scores on the Stanford Sleepiness Scale (SSS) at each test session for Days 1 and 2.

| Measures | RT (log) |  |  | SSS scores |  |  |
| --- | --- | --- | --- | --- | --- | --- |
|  | pretraining | Posttraining | postsleep | pretraining | Posttraining | Postsleep |
| Day 1 | 2.52 ± 0.01 | 2.52 ± 0.01 | 2.49 ± 0.01 | 2 (1-2) | 2 (1-3) | 1 (1-2) |
| Day 2 | 2.51 ± 0.01 | 2.51 ± 0.01 | 2.51 ± 0.01 | 2 (1-2) | 2 (1-2) | 1 (1-2) |
| Z values | -1.396 | -1.207 | -1.491 | 0.000 | -2.449 | 0.000 |
| P values | 0.163 | 0.227 | 0.136 | 1.000 | 0.014 | 1.000 |

The RTs for PVT are presented as the mean ± SEM, and the SSS scores are presented as the median and range, along with the results of the Wilcoxon signed-rank test (n=17).

Next, we examined the sleep duration the night before each experiment using a sleep log. The participants slept for 7.9 ± 0.2 hours (mean ± SEM) at home before Day 1 and for 7.8 ± 0.3 hours (mean ± SEM) before Day 2. There was no significant difference in sleep duration before the experiments on Days 1 or 2 (paired t test, t(16)=-0.237, p=0.816, 95% CI = [-0.2 0.1]). These results suggest that the smaller offline performance gain on Day 1 was not caused by a shorter sleep duration before the experiment.

Finally, we tested whether the impact of sleep disturbance was associated with the degree of decline in the offline performance gains. The difference in the SOL between Day 2 and Day 1 was used to indicate the degree of sleep disturbance. Similarly, the difference in the amount of offline performance gain between Days 1 and 2 would indicate the degree of decline in behavioral visual plasticity. The correlation coefficients between them were not significantly correlated (r=0.005, p=0.984).

Thus, the results of Experiment 1 showed that offline performance gains that occur during a normal sleep session were significantly attenuated by the FNE among young, healthy participants. In addition, the results suggest that the significantly larger offline performance gains on Day 2 were not caused by learning transfer. Moreover, the smaller offline performance gains on Day 1 were explained by sleepiness or by the sleep duration the night before the experiment. These results consistent with the hypothesis that the FNE disrupts visual plasticity.

### Experiment 2

In the next experiment, we tested whether the FNE impairs the neurochemical processing involved in visual plasticity during NREM sleep. We compared the E/I ratios, an index of visual plasticity ^7,27-29^, during NREM sleep on Day 1 and Day 2. A new group of participants (n=15) slept for approximately 1 hour twice (Day 1 and Day 2) in the MRI scanner together with a PSG setting (see methods). MRS was continuously performed during the session to measure the concentrations of excitatory (Glx, a combination of glutamate and glutamine) and inhibitory (GABA) neurotransmitters in early visual areas. The E/I ratio was calculated as the concentration of Glx divided by that of GABA during wakefulness and NREM sleep, respectively ^30^, for each participant. Note that part of the E/I ratio data was published in a previous study ^7^ and that new analyses were conducted using the same dataset.

First, we tested whether the FNE occurred during the first sleep session (**Table 3**). As in Experiment 1, we calculated the SOL according to previous studies ^16^ to quantify the FNE. The SOL was significantly different between the Day 1 session and Day 2 session (**Fig. 2A**; *Z*=2.54, *p*=0.011, 95% CI=[2.7, 14.8]). The SOL was significantly longer on Day 1 than on Day 2, supporting the FNE on Day 1.

**Fig. 2.**
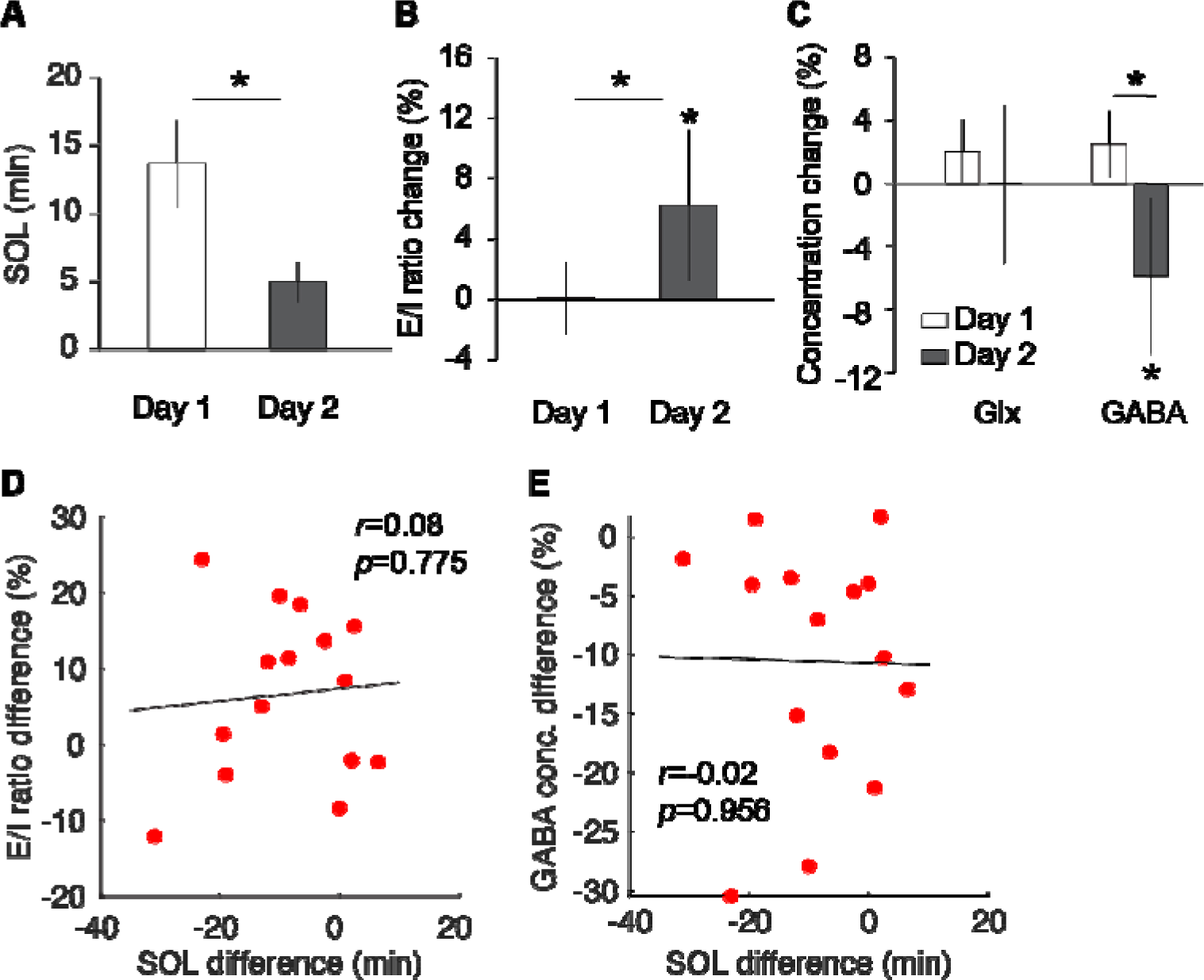
Results of Experiment 2. **A.** Sleep onset latency (SOL). **B.** The E/I ratio changes during NREM sleep compared to wakefulness. **C.** Glx and GABA concentration changes compared to wakefulness. Paired or one-sample t test, *p<.05. **D.** Scatterplot between the sleep-onset latency (SOL) difference and E/I ratio change difference (Day 2 - 1). **E.** Scatter plot between the sleep-onset latency (SOL) difference and GABA concentration (conc.) difference (Day 2 - 1). N=15.

**Table 3:** Sleep parameters for Experiment 2.

|  | Day 1 |  | Day 2 |  | Wilcoxon signed rank test |  |
| --- | --- | --- | --- | --- | --- | --- |
|  |  |  |  |  | Z | P |
| SOL (min) | 13.8 | ± 3.16 | 4.9 | ± 1.51 | 2.54 | 0.011 |
| WASO (min) | 8.5 | ± 1.67 | 12.0 | ± 3.06 | -0.85 | 0.394 |
| Stage W (%) | 24.9 | ± 4.05 | 21.8 | ± 4.34 | 0.80 | 0.427 |
| Stage N1 (%) | 32.3 | ± 5.02 | 15.6 | ± 2.34 | 2.90 | 0.004 |
| Stage N2 (%) | 34.8 | ± 4.40 | 47.8 | ± 5.34 | -2.39 | 0.017 |
| Stage N3 (%) | 13.6 | ± 3.98 | 21.0 | ± 6.24 | -2.20 | 0.028 |
| Stage R (%) | 7.8 | ± 3.08 | 7.8 | ± 2.42 | 0.45 | 0.655 |
| Total time (min) | 63.1 | ± 4.47 | 67.3 | ± 4.44 | -0.60 | 0.551 |

SOL, sleep-onset latency. WASO, wake after sleep onset. SE, sleep efficiency. Each sleep stage duration was measured from the first sleep cycle. The total time was defined as the sum of W, N1, N2, N3 and R stage durations within the first sleep cycle. Values are means ± SEs. Wilcoxon signed-rank test was applied to test whether sleep parameters were significantly different between sleep sessions (days 1 vs. 2, n=15).

We next tested whether the FNE affected the change in the E/I ratio during sleep. We calculated the mean change in the E/I ratio during NREM sleep relative to baseline, which was the average E/I ratio during wakefulness, for each sleep session. We tested whether the mean E/I ratio during NREM sleep was significantly different between Day 1 (when the FNE occurred) and Day 2 (when the FNE subsided).

As we reported before, the mean E/I ratio during NREM sleep on Day 2, during which natural sleep progresses, was significantly greater than that at baseline (**Fig. 2B**; one sample t test against 0, t(14)=2.95, p=0.011, 95% CI =[1.7, 10.7]) ^7^. However, on Day 1, when the FNE occurred, the mean E/I ratio change during NREM sleep was not significantly different from zero (one sample t test against 0, t(14)=-0.22, p=0.830, 95% CI =[-5.3, 4.3]). Thus, the mean E/I ratio during NREM sleep relative to baseline was significantly different between the Day 1 and Day 2 sleep sessions (paired t test, t(14)=- 2.38, p=0.032, 95% CI=[-12.8, -0.7]). The results suggest that FNE affects the change in the E/I ratio during NREM sleep, preventing visual plasticity from increasing during sleep.

A previous study showed that the GABA concentration, not the Glx concentration, decreased during NREM sleep, which resulted in an increased E/I ratio ^7^. Thus, we tested whether the GABA or Glx concentration during NREM sleep was affected by the FNE (**Fig. 2C**). We calculated the mean changes in GABA and Glx concentrations during NREM sleep relative to baseline, which was the average amount during wakefulness. The magnitude of change in the GABA concentration was significantly lower on Day 2 than on Day 1 (paired t test, t(14)=4.00, p=0.001, 95% CI=[4.9, 16.2]). Furthermore, the magnitude of change in the GABA concentration was significantly different from zero on Day 2 (one-sample t test against 0, t(14)=-3.98, p=0.001, 95% CI=[-9.02, -2.70]) but not on Day 1 (one-sample t test against 0, t(14)=1.8, p=0.086, 95% CI=[-0.8, 10.1]). In contrast, there was no significant difference in the mean change in the Glx concentration between Days 1 and 2 during NREM sleep (paired t test, t(14)=1.85, p=0.085, 95% CI=[-0.5, 7.1]). The Glx concentration was not significantly different from zero for either Day 1 (one-sample t test against 0, t(14)=1.72, p=0.108, 95% CI=[-0.8, 7.3]) or Day 2 (one-sample t test against 0, t(14)=-0.02, p=0.983, 95% CI=[-3.6, 3.5]), suggesting that the Glx concentration was not affected by the FNE. These results showed that the GABA concentration, but not the Glx concentration, is affected by the FNE.

We next tested whether the impact of sleep disturbance induced by the FNE was correlated with a change in neurochemical processing during sleep. We measured the difference in the SOL and E/I ratio and GABA concentration between days. We examined the correlation between the SOL difference (Day 2 - Day 1) and the E/I ratio difference (Day 2 - Day 1) as well as the correlation between the SOL difference (Day 2 - Day 1) and the GABA concentration difference (Day 2 - Day 1). However, we did not find any associations between the SOL and the change in the mean E/I ratio (r=0.08, p=0.775; CI=[- 0.5, 0.6]; **Fig. 2D**) or between the SOL and the change in the GABA concentration (r=-0.02, p=0.956; CI=[-0.5, 0.5]; **Fig. 2E**). The results suggest that processes associated with the modulation of vigilance, which induces the FNE, and the neural processes inducing neurochemical visual plasticity during sleep are independent of each other.

Additional control analyses indicated that the significant difference in the E/I ratio change between the two sessions could not be attributed to factors such as MRS data quality, sleepiness before the nap, or sleep duration prior to the recorded sleep session. First, we tested whether the quality of the MRS data differed between the two sessions (**Table 4**). However, neither the shimming, %SD, NAA linewidth, nor frequency drift measures were significantly different between the two sessions (**Table 4**). Second, a nonparametric test showed that the median SSS score ^32^, a subjective sleepiness measure, was not significantly different between the two sessions (median = 2 and range = 1-3 on both Days 1 and 2; Wilcoxon signed-rank test, Z = 1.34, p=0.180, n=9). Third, the average length of sleep at home the night before the experiments was not significantly different between the two sessions (on both Days 1 and 2, 7.9 ± 0.2; paired t test, t(14) = 0.02, p=0.982). Because these data were not significantly different between Days 1 and 2, the significant difference in the relative E/I ratio change during NREM sleep between days is not attributable to these factors. These results suggest that any factors other than the FNE account for the significant differences in neurochemical processing during sleep between Days 1 and 2.

**Table 4.**
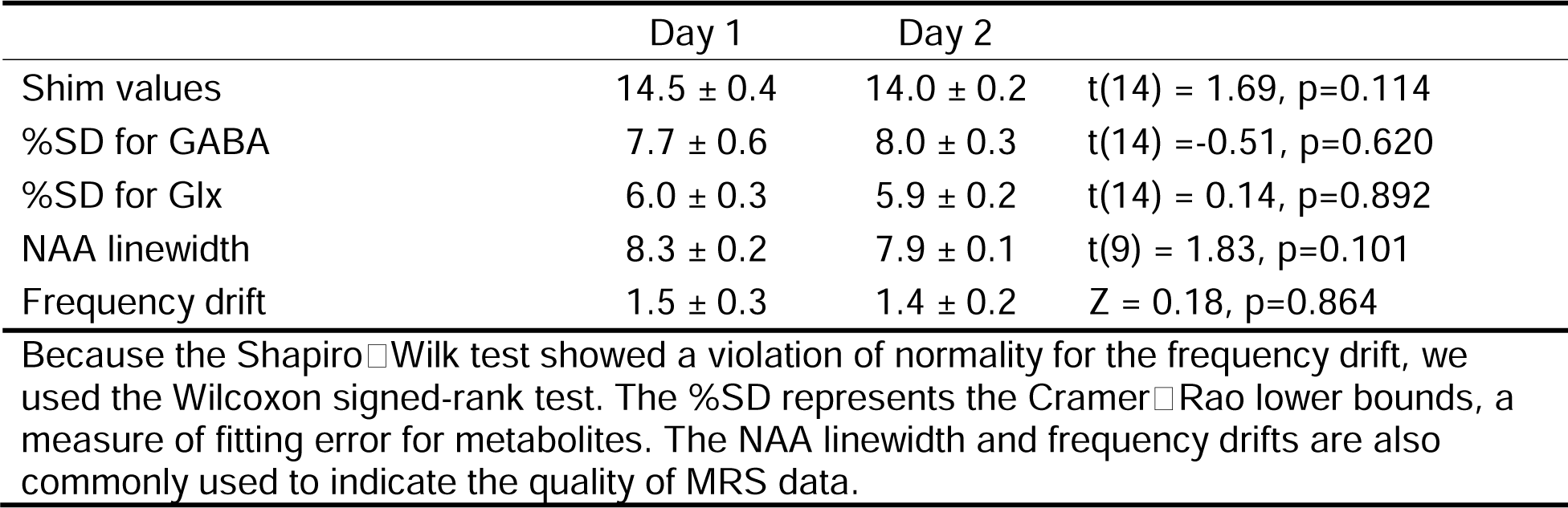
MRS data quality on Day 1 and 2 in Experiment 2.

| **Table 4.** MRS data quality on Day 1 and 2 in Experiment 2. |  |  |  |
| --- | --- | --- | --- |
|  | Day 1 | Day 2 |  |
| Shim values | $14.5 \pm 0.4$ | $14.0 \pm 0.2$ | $t(14) = 1.69$, $p=0.114$ |
| %SD for GABA | $7.7 \pm 0.6$ | $8.0 \pm 0.3$ | $t(14) = -0.51$, $p=0.620$ |
| %SD for Glx | $6.0 \pm 0.3$ | $5.9 \pm 0.2$ | $t(14) = 0.14$, $p=0.892$ |
| NAA linewidth | $8.3 \pm 0.2$ | $7.9 \pm 0.1$ | $t(9) = 1.83$, $p=0.101$ |
| Frequency drift | $1.5 \pm 0.3$ | $1.4 \pm 0.2$ | $Z = 0.18$, $p=0.864$ |
| Because the Shapiro-Wilk test showed a violation of normality for the frequency drift, we used the Wilcoxon signed-rank test. The %SD represents the Cramer-Rao lower bounds, a measure of fitting error for metabolites. The NAA linewidth and frequency drifts are also commonly used to indicate the quality of MRS data. |  |  |  |

Together, the results of Experiments 1 and 2 are consistent with the hypothesis that the FNE disrupts visual plasticity during sleep. The results further suggest that the degree of the FNE and the index of behavioral visual plasticity during sleep are independent of each other.

## Discussion

The present study investigated whether and how the FNE affects visual plasticity during sleep among healthy young individuals. The results clearly showed that even temporary sleep disturbances in healthy participants affect visual plasticity both behaviorally (Experiment 1) and neurochemically (Experiment 2). Additionally, several control analyses suggest that the significant changes in performance and neurochemical processing associated with the FNE were not caused by potentially confounding factors, such as interactions between learning sessions on two days, sleepiness, sleep duration before the experiment, or MRS data quality.

Interestingly, SOL, an indicator of the degree of the FNE, was not significantly correlated with the amount of offline performance gains during the sleep session (Experiment 1); furthermore, SOL was not significantly correlated with an increase in the E/I ratio or a decrease in the GABA concentration during sleep (Experiment 2). These results suggest that the degree of sleep disturbance and the degree of visual plasticity are not directly associated when sleep disturbances alter neurochemical processing in early visual areas or impair visual plasticity in healthy participants.

Why does the FNE impair visual plasticity during sleep when there is no direct correlation between the processes involved in the FNE and visual plasticity outcomes (i.e., the E/I ratio and offline performance gains)? We propose that there is an indirect link between these processes, even though these two processes occur separately (**Fig. 3**). To discuss this possibility, we first describe the process involved in the FNE, followed by the process of visual plasticity. Finally, we discuss the possible indirect links between them.

**Fig. 3.**
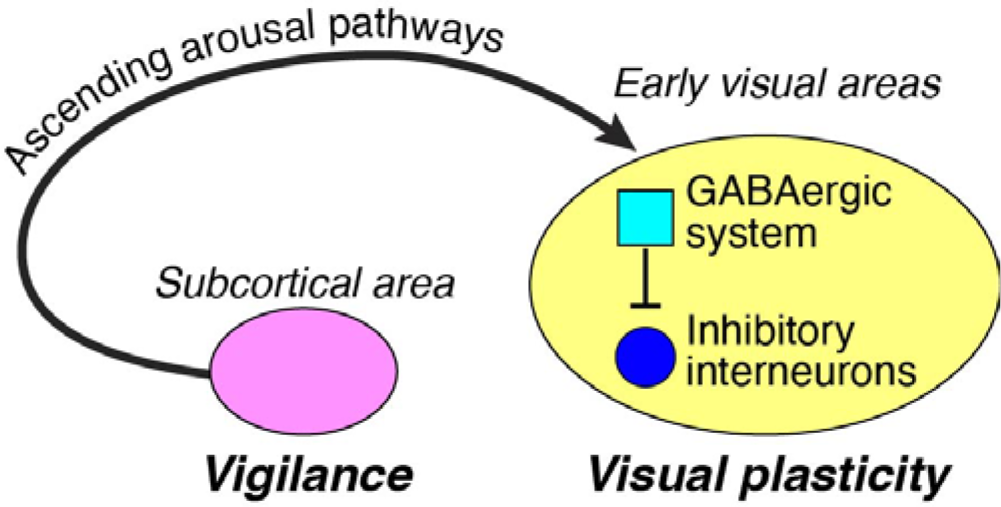
The proposed model for the interaction between the FNE and the visual plasticity process. The FNE is associated with increased vigilance to monitor the novel surroundings, whereas visual plasticity is involved in the disinhibition of inhibitory interneurons. The FNE may impact visual plasticity through ascending arousal pathways, which are not suppressed enough during disturbed sleep.

Previous studies have shown that the FNE involves increased vigilance during sleep to monitor novel surroundings ^16^. Increased vigilance, which is associated with the FNE, is evident through several indicators: reduced slow wave activity in brain regions linked to the default mode network, an increase in the number of arousals, and amplified amplitudes of event-related potentials during sleep, all of which are known to correlate with vigilance ^16,19^. This increased vigilance is likely to be governed by sleep-wake regulators in subcortical areas.

On the other hand, during undisturbed human sleep, early visual areas are implicated in enhanced visual plasticity, as demonstrated by the increases in the E/I ratio attributed to decreased GABA concentrations ^7^. Notably, another line of research showed that GABAergic inhibitory interneurons in early visual areas are involved in visual plasticity ^33,34^. If interneurons increase their activity to enhance plasticity, the GABA concentration should increase in early visual areas. However, the GABA concentration in early visual areas was decreased during undisturbed sleep in humans ^7^. This finding suggested that another GABAergic system exists and decreases its activity during undisturbed sleep. Perhaps the additional GABAergic system decreases its activity during undisturbed sleep, thus causing the inhibitory interneurons to be disinhibited enhancing visual plasticity ^7^.

Based on the processing of the FNE and visual plasticity, we hypothesize that there is a link between the FNE and visual plasticity, i.e., a connection between subcortical neurons involved in regulating sleep-wake cycles and neurons in early visual areas. These connections potentially involve ascending arousal pathways, which are responsible for vigilance and originate from subcortical neurons projecting to the cerebral cortex via various key neurotransmitters to stimulate brain activity ^35^.

Since the ascending arousal pathways promote wakefulness, they need to be shut off during sleep. A failure to adequately shut off the pathways likely causes sleep problems. Previous studies have shown that lesions in these pathways lead to prolonged sleep ^35^, whereas increased activation of these pathways is observed during sleep in insomniac patients ^36^.

As mentioned above, our proposed model assumes the existence of two separate forms of GABAergic systems in early visual areas. The first GABAergic system involves inhibitory interneurons (blue circle in **Fig. 3**), which are disinhibited from one another (cyan square in **Fig. 3**) during sleep. The second GABAergic system (cyan square in **Fig. 3**) is sensitive to and influenced by ascending arousal pathways, which could remain active during sleep in the presence of the FNE. The active ascending arousal pathways might activate the second GABAergic system in early visual areas and prevent the interneurons from being adequately disinhibited during sleep with the FNE, leading to disrupted visual plasticity. The results of Experiment 2 showing that the GABA concentrations did not decrease during sleep with the FNE, are in accordance with the proposed FNE model and impaired visual plasticity.

This study has several limitations. First, the present study used the FNE as a model for disturbed sleep. Visual plasticity is altered by chronic sleep disturbances ^37^ or sleep restrictions ^38^. While it is possible that a comparable mechanism is shared between chronic sleep problems and the FNE, it remains unclear how much the FNE could be generalized to other chronic sleep problems.

Second, we focused on the analysis of NREM sleep. However, both NREM sleep and REM sleep are involved in facilitating learning ^1,22^. Previous studies have demonstrated that REM sleep is critical for stabilizing presleep learning, thus making learning robust against interference, and for facilitating new postsleep learning ^7,39^. Due to the technical limitations pertaining to MRI scanning and our protocols, we were only able to measure brain activity for a maximum of 90 minutes of daytime sleep. With this time limitation, it is difficult to acquire enough REM sleep data, specifically on Day 1, since the latency to REM sleep is prolonged, and the first bout of REM sleep is occasionally skipped due to the FNE ^40^. Thus, limited REM sleep data were obtained in the present study. Our previous study suggested that REM sleep is also modulated by the FNE ^19^. Thus, future studies need to investigate whether and how the FNE affects the learning-facilitation process during REM sleep.

Third, how the FNE influences other types of learning and memory needs to be clarified. The role of sleep in learning is not limited to VPL; other domains of learning and memory, including declarative memory ^41-45^ and motor learning ^5,46-49,^ have been widely studied. The present study focused solely on the neurochemical process in early visual areas associated with VPL. Future studies need to investigate whether and, if so, how sleep disturbances affect various forms of learning and memory, in addition to their impact on neurochemical processing in brain regions associated with these types of learning and memory.

Finally, in the present study, healthy young participants with a regular sleep-wake schedule and no self-reported chronic sleep problems were recruited. However, how aging affects the interaction between the FNE and brain plasticity during sleep has yet to be determined. The number of individuals aged 65 years or older is increasing ^8,50^. The aging of society will inevitably result in an increasing number of individuals with chronic sleep problems, as aging is associated with an increasing prevalence of various factors affecting sleep, including medical and psychiatric comorbidities, medication use, and primary sleep disorders ^51-54^. Future research needs to clarify how various aging-related sleep disturbances, including difficulty falling asleep, frequent awakenings, and early morning awakenings, influence brain plasticity during sleep.

## Acknowledgments

This work was supported by the NIH (R01EY031705, R01EY019466, R01EY027841). Part of this research was also supported by the JSPS KAKENHI grant numbers JP20K22297, JP22H01107, and JP22K18664; the Naito Foundation; the Takeda Science Foundation; and the Uehara Foundation.

## Author Contributions

MT and YS designed the study. MT, TBD, and ZW performed the experiments. MT, TY, TBD, and ZW analyzed the data. MT, TW, and YS wrote the manuscript.

## Methods

Here, we describe the procedures common to the two experiments, followed by the procedures specific to each experiment.

## 1. Procedures common to the two experiments

### 1.1. Participants

There were criteria for eligibility for participation in the present study. First, we excluded people who frequently played action video games because extensive video gaming affects visual and attention processing ^10,55,56^. Second, we excluded people with prior experience in visual perceptual learning tasks because previous VPL experiences may alter visual processing ^21,57^. Third, participants needed to have normal or corrected-to-normal vision. Fourth, we excluded people with an irregular sleep schedule. Fifth, we excluded people who had a physical or psychiatric disease, were currently taking medication, or were suspected of having a sleep disorder based on self-report ^16,18,58^. Finally, we recruited young adults who were older than 18 years old but excluded individuals older than 30 due to known altered sleep structures ^59^.

After careful screening, 36 young adults participated in the study. Nineteen participants participated in Experiment 1. Two participants from Experiment 1 were excluded from further analyses due to irregular sleep-wake habits detected from actigraphy data and a sleep log. The remaining 17 participants’ data were included in Experiment 1 (7 females and 10 males, 23.4 ± 0.84 years old, mean ± SEM). A different group of 17 participants participated in Experiment 2. Two participants were excluded from further analyses since they showed no wakefulness lasting at least 2 minutes, which was necessary to normalize of MRS data (see **3.2** below). The remaining 15 participants were included in Experiment 2 (7 females and 8 males, 24.2 ± 1.06 years old; mean ± SEM). The data for Day 2 in Experiment 2 in the present study were also used in part of a previous study ^7^.

All the participants provided written informed consent to participate in the experiments. This study was approved by the institutional review board of Brown University.

### 1.2. Experimental design

Participants in both experiments underwent two sessions, designated Day 1 and Day 2. These sessions were conducted approximately one week apart so that any effects of disrupted sleep due to the FNE on Day 1 would not carry over to the second sleep session.

Starting 3-7 days before the onset of the sleep sessions, the participants were instructed to maintain their regular sleep-wake habits, i.e., their daily wake/sleep time and sleep duration. On the day before the sleep session, they were instructed to refrain from consuming alcohol, engaging in unusually excessive physical exercise, and taking naps. Their sleep-wake habits were monitored by an actigraph and a sleep log. Caffeine consumption was not allowed on the days of the experiments. Several questionnaires related to sleep quality and habits were administered prior to the Day 1 session, including the Pittsburgh Sleep Quality Index questionnaire ^60^, the Morningness-Eveningness Questionnaire ^61^ and the Edinburgh Handedness Questionnaire ^62^ (data not shown).

The sleep sessions started in the early afternoon (approximately 1-3 pm) each day after the electrodes were attached for PSG measurement (see **1.3** below). The sleep sessions were conducted as follows. For Experiment 1, the sleep session was conducted in the MRI scanning room. MRI, MRS, and PSG data were measured simultaneously during sleep (see **3.1** below and elsewhere ^30^ for details). The total duration of sleep varied between 45 and 90 minutes. For Experiment 2, the sleep session was recorded via PSG (see **1.3**) in an electrically shielded and sound-attenuated sleep chamber. The total duration of sleep was 90 min. After each sleep session, the participants answered a questionnaire about their sleep during the session (data not shown).

### 1.3. PSG recording

The preparation for PSG took approximately 30 min. All the data were recorded with a standard amplifier (BrainAmp MR or BrainAmpStandard, Brain Products) and software (BrainVision Recorder, Brain Products). PSG consisted of electroencephalography (EEG), electrooculography (EOG), and electromyography (EMG) data. Electrocardiograms (ECGs) were also examined for Experiment 1. EEG data were recorded at 23-31 scalp sites according to the 10% system of electrode positioning. EOG data were recorded from electrodes placed at the outer canthi of both eyes (horizontal EOG) and above and below the left and right eyes (vertical EOG). EMG data were recorded from electrodes placed at the mentum. ECGs were recorded from the lower shoulder blade. The sampling frequency ranged from 500-5000 Hz. The online reference was Fz, and the measurements were rereferenced to the average of the left (TP9) and right (TP10) mastoids after the recording was complete. In Experiment 1, since PSG was collected simultaneously with MRS, the PSG was obscured by the MRI scanner and by ballistocardiogram artifacts. This noise was removed using Brain Vision Analyzer 2 (Brain Products, LLC) before sleep-stage scoring (see **1.4** for sleep-stage scoring).

### 1.4. Sleep-stage scoring and sleep parameters

Sleep stages were scored in 30-s epochs using the standard criteria stated by the American Academy of Sleep Medicine (AASM) Scoring Manual ^63^. The stages were as follows: wakefulness (stage W), NREM stage 1 sleep (stage N1), NREM stage 2 sleep (stage N2), NREM stage 3 sleep (stage N3), and REM sleep. Standard sleep parameters, including sleep-onset latency (SOL, the latency to the first appearance of stage N2 after the lights were turned off), were obtained to indicate the general sleep structure for each experiment ^64^).

### 1.5. Sleepiness measurement

Sleepiness was measured by the subjective Stanford Sleepiness Scale (SSS) ^32^ with (Experiment 1) or without (Experiment 2) a psychomotor vigilance test (PVT) ^31^.

The SSS scores ranged from 1 (feeling active, vital, alert, or wide awake) to 7 (no longer fighting sleep, sleep onset soon; having dream-like thoughts); participants chose the score that corresponded to their state of sleepiness. The PVT was implemented with open-source Psychology Experiment Building Language (PEBL) software ^31^. In each trial of the PVT, after a fixation screen was displayed, a target screen was presented in which a red circle appeared in the center of the screen. The participants were required to press the spacebar on a keyboard as quickly as possible upon detection of the circle. The time interval between the fixation screen and the screen with a red circle varied between 1000 and 4000 ms. The task lasted approximately 2 min ^65,66^. The reaction time data were measured and log-transformed to reduce skewness. The average reaction time was obtained as a measurement of behavioral sleepiness.

In Experiment 1, in addition to the SSS, the participants performed a PVT to measure behavioral sleepiness prior to each test session. In Experiment 2, 9 participants rated their sleepiness before PSG preparations using the SSS.

### 1.6. Statistical analyses

An α level (type I error rate) of 0.05 was set for all the statistical analyses, and two-tailed tests were used. The ShapiroLJWilk test was conducted for all the data to test whether the data were normally distributed. Nonparametric tests (the MannLJWhitney U test or the Wilcoxon signed-rank test) were used if the normality test rejected the null hypothesis. Levene’s test was used to test for homogeneity of variance before ANOVA. It was confirmed that homogeneity of variance was not violated in any of the data intended for the ANOVA (all *p* > 0.05). A Grubbs test was conducted to detect outliers. When a t test was used, we also computed the 95% confidence interval (CI). To analyze performance improvement, ANOVA was first conducted, and then t tests were conducted as post hoc tests. All the statistical tests were conducted in SPSS (ver. 22, IBM Corp.) and MATLAB (R2020a, The MathWorks, Inc.).

## 2. Procedures specific to Experiment 1

### 2.1. Texture discrimination task

We used the texture discrimination task (TDT) ^57^, a standard VPL task previously shown to be facilitated by sleep ^22,26,67,68,^ in Experiment 1. The TDT consists of two tasks: the orientation task and the letter task. The orientation task was the main task, whereas the letter task was designed to control participants’ eye fixation ^57^. The orientation task shows retinotopic location specificity, where learning occurs only in the trained visual field location ^57,69^. In addition, learning is specific to the orientation of background lines ^57,70^. These findings suggest that the early visual areas in which low-level visual features are processed are critically involved in the improvement of TDT, as supported by neuroimaging studies ^4,23,25,69,71^.

The target location for the orientation task was either the left or the right side of the visual hemifield within a 7–9° eccentricity. The visual hemifield for the target location was changed from Day 1 to Day 2 so that learning each day would be independent owing to the location specificity of learning in the TDT ^57^. For 9 participants, the target location was the left hemifield on Day 1 and the right hemifield on Day 2. The target location for the remaining 8 participants was the right hemifield on Day 1 and the left hemifield on Day 2.

The time interval between the target onset and mask was referred to as the stimulus-to-mask onset asynchrony (SOA). The SOA was modulated across trials to control for task difficulty. When the mask stimulus is presented, new information processing in the retina overrides information about the target stimulus. Thus, the task difficulty increases with decreasing SOA. We describe the summary of the procedure below (the TDT has been described previously ^6,24^).

Each test session took 6-10 min. There were 5-6 SOAs, ranging between 33 and 316 ms. The total number of trials ranged from 75 to 120 trials (15-20 trials per SOA). The presentation order of the SOAs was pseudorandomized to reduce the amount of learning and fatigue during test sessions ^72^. The same SOAs were used in the training sessions as in the test sessions. The total number of trials in the training session was 600-720. The training session started with the longest SOA (300 ms) and proceeded in descending order.

The stimuli used were generated by MATLAB software with Psychtoolbox (http://psychtoolbox.org) ^73,74^.

### 2.2. TDT performance

The TDT performance was measured based on the threshold SOAs in milliseconds corresponding to 80% correct performance as follows. First, the percentage of correct responses for the orientation task was obtained for each SOA. Second, a cumulative Gaussian function was fitted for each psychometric data point to determine the threshold for each subject for each test session using the psignifit toolbox (ver. 2.5.6) for MATLAB ^75^. All trials were included for the estimation of threshold SOA except for trials in which the letter task was incorrect, as incorrect trials for the letter task suggest that participants did not effectively maintain eye fixation during these trials. Finally, TDT performance changes were computed based on relative changes in the threshold SOAs (ms) between test sessions. The change in performance (%) with training was calculated as [100 × (pretraining – posttraining)/(pretraining)]. Similarly, the change in performance from posttraining to posttraining (%) corresponding to offline performance gains by sleep was calculated as [100 × (posttraining – postsleep)/(posttraining)].

## 3. Procedures specific to Experiment 2

### 3.1. Anatomical MRI and MRS acquisition

Anatomical MRI and MRS data were collected in Experiment 2. Refer to our previous research ^7,30^ for details regarding simultaneous MRS and PSG scans and how we coregistered MRS data to sleep stage data.

Participants were scanned using a 3T Siemens Prisma scanner with a 64-channel head coil. For structural MRI, T1-weighted MR images (MPRAGE; 256 slices, voxel size = 1 mm^3^, 0 mm slice gap) were collected. For five participants, we ran the MEGA-PRESS and PRESS scans alternately as an exploratory method during the sleep session. We used the data that were obtained only from the MEGA-PRESS scans so that the neurotransmitter concentrations would be estimated from the same type of sequence ^7^. The acquisition time for the MEGA-PRESS scans with water suppression was 3.3 min (TR = 1.5 s, TE = 68 ms, number averaged = 64, VOI = 2×2×2 cm^3^, scan time 198 s including 6-s dummy scans for the steady state of longitudinal magnetization). For the remaining ten participants, we ran only the MEGA-PRESS scans during the sleep session since we found that using a consistent scan was more efficient than alternating two different scans ^76,77^ (TR = 1.25 s, TE = 68 ms, number averaged = 240, VOI = 2.2×2.2×2.2 cm^3^) with double-banded pulses used to simultaneously suppress the water signal and edit the γ-CH2 resonance of GABA at 3.0 ppm ^7^. The volume of interest (VOI) was early visual areas (determined anatomically on the calcarine sulcus).

## 3.2 MRS analysis

Spectroscopic imaging data were processed using LCModel ^78,79^ for metabolite quantification, including Glx, GABA, and NAA. The LCModel assumes that the obtained spectrum can be fitted in the frequency domain using a linear combination of basis functions. To quantify the Glx and GABA concentrations, we divided the raw Glx and GABA concentrations by the NAA concentration.

The E/I ratio was calculated as the concentration of Glx divided by the concentration of GABA for each sleep stage. The average E/I ratio during wakefulness was regarded as the baseline E/I ratio ^7,30^. The relative change (%) in the E/I ratio during NREM sleep was calculated as [(E/I_NREMsleep_ – E/I_wake_)/E/I_wake_ × 100]. We also calculated relative changes (%) in Glx and GABA concentrations during NREM sleep compared to those during wakefulness in a similar manner to the calculation of relative changes in the E/I ratio.

To measure the MRS data quality, we obtained the CramerLJRao lower bounds for GABA and Glx, the shim values, the frequency of the NAA signal (or NAA linewidth), and the frequency drift. The reliability of the quantification of Glx and GABA was indicated by the CramerLJRao lower bounds (or %SD) as a measure of fitting errors obtained by the LCModel. The NAA linewidth, where larger values suggest more head motion, was also noted for each measurement. Since the raw data (twix files) were obtained from 10 out of 15 participants, the NAA linewidth and frequency drift, which require twix files, were measured from these 10 participants.

## References

1. Mednick, S., Nakayama, K., and Stickgold, R. (2003). Sleep-dependent learning: a nap is as good as a night. Nat Neurosci 6, 697–698. 10.1038/nn1078.

2. Schönauer, M., Geisler, T., and Gais, S. (2014). Strengthening procedural memories by reactivation in sleep. J Cogn Neurosci 26, 143–153. 10.1162/jocn_a_00471.

3. Spencer, R.M., Sunm, M., and Ivry, R.B. (2006). Sleep-dependent consolidation of contextual learning. Curr Biol 16, 1001–1005. 10.1016/j.cub.2006.03.094.

4. Stickgold, R., James, L., and Hobson, J.A. (2000). Visual discrimination learning requires sleep after training. Nat Neurosci 3, 1237–1238. 10.1038/81756.

5. Tamaki, M., Huang, T.R., Yotsumoto, Y., Hämäläinen, M., Lin, F.H., Náñez, J.E., Sr., Watanabe, T., and Sasaki, Y. (2013). Enhanced spontaneous oscillations in the supplementary motor area are associated with sleep-dependent offline learning of finger-tapping motor-sequence task. J Neurosci 33, 13894–13902. 10.1523/jneurosci.1198-13.2013.

6. Tamaki, M., and Sasaki, Y. (2022). Sleep-Dependent Facilitation of Visual Perceptual Learning Is Consistent with a Learning-Dependent Model. J Neurosci 42, 1777–1790. 10.1523/jneurosci.0982-21.2021.

7. Tamaki, M., Wang, Z., Barnes-Diana, T., Guo, D., Berard, A.V., Walsh, E., Watanabe, T., and Sasaki, Y. (2020). Complementary contributions of non-REM and REM sleep to visual learning. Nat Neurosci 23, 1150–1156. 10.1038/s41593-020-0666-y.

8. Institute of Medicine Committee on Sleep, M., and Research (2006). The National Academies Collection: Reports funded by National Institutes of Health. In Sleep Disorders and Sleep Deprivation: An Unmet Public Health Problem, H.R. Colten, and B.M. Altevogt, eds. (National Academies Press (US). Copyright © 2006, National Academy of Sciences.). 10.17226/11617.

9. Léger, D., and Bayon, V. (2010). Societal costs of insomnia. Sleep Med Rev 14, 379–389. 10.1016/j.smrv.2010.01.003.

10. Green, C.S., and Bavelier, D. (2003). Action video game modifies visual selective attention. Nature 423, 534–537. 10.1038/nature01647.

11. Lie, J.D., Tu, K.N., Shen, D.D., and Wong, B.M. (2015). Pharmacological Treatment of Insomnia. P t 40, 759–771.

12. Madari, S., Golebiowski, R., Mansukhani, M.P., and Kolla, B.P. (2021). Pharmacological Management of Insomnia. Neurotherapeutics 18, 44–52. 10.1007/s13311-021-01010-z.

13. Agnew, H.W., Jr., Webb, W.W., and Williams, R.L. (1967). Sleep patterns in late middle age males: an EEG study. Electroencephalogr Clin Neurophysiol 23, 168–171. 10.1016/0013-4694(67)90107-1.

14. Rechtschaffen, A., and Verdone, P. (1964). AMOUNT OF DREAMING: EFFECT OF INCENTIVE, ADAPTATION TO LABORATORY, AND INDIVIDUAL DIFFERENCES. Percept Mot Skills 19, 947–958. 10.2466/pms.1964.19.3.947.

15. Ding, L., Chen, B., Dai, Y., and Li, Y. (2022). A meta-analysis of the first-night effect in healthy individuals for the full age spectrum. Sleep Med 89, 159–165. 10.1016/j.sleep.2021.12.007.

16. Tamaki, M., Bang, J.W., Watanabe, T., and Sasaki, Y. (2016). Night Watch in One Brain Hemisphere during Sleep Associated with the First-Night Effect in Humans. Curr Biol 26, 1190–1194. 10.1016/j.cub.2016.02.063.

17. Tamaki, M., Nittono, H., and Hori, T. (2005). The first-night effect occurs at the sleep-onset period regardless of the temporal anxiety level in healthy students. Sleep and Biological Rhythms 3, 92–94. 10.1111/j.1479-8425.2005.00167.x.

18. Tamaki, M., Bang, J.W., Watanabe, T., and Sasaki, Y. (2014). The first-night effect suppresses the strength of slow-wave activity originating in the visual areas during sleep. Vision Res 99, 154–161. 10.1016/j.visres.2013.10.023.

19. Tamaki, M., and Sasaki, Y. (2019). Surveillance During REM Sleep for the First-Night Effect. Front Neurosci 13, 1161. 10.3389/fnins.2019.01161.

20. Dosher, B.A., and Lu, Z.L. (2009). Hebbian Reweighting on Stable Representations in Perceptual Learning. Learn Percept 1, 37–58. 10.1556/lp.1.2009.1.4.

21. Watanabe, T., Náñez, J.E., and Sasaki, Y. (2001). Perceptual learning without perception. Nature 413, 844–848. 10.1038/35101601.

22. Karni, A., Tanne, D., Rubenstein, B.S., Askenasy, J.J., and Sagi, D. (1994). Dependence on REM sleep of overnight improvement of a perceptual skill. Science 265, 679–682. 10.1126/science.8036518.

23. Schwartz, S., Maquet, P., and Frith, C. (2002). Neural correlates of perceptual learning: a functional MRI study of visual texture discrimination. Proc Natl Acad Sci U S A 99, 17137–17142. 10.1073/pnas.242414599.

24. Tamaki, M., Wang, Z., Watanabe, T., and Sasaki, Y. (2019). Trained-feature-specific offline learning by sleep in an orientation detection task. J Vis 19, 12. 10.1167/19.12.12.

25. Walker, M.P., Stickgold, R., Jolesz, F.A., and Yoo, S.S. (2005). The functional anatomy of sleep-dependent visual skill learning. Cereb Cortex 15, 1666–1675. 10.1093/cercor/bhi043.

26. Yotsumoto, Y., Sasaki, Y., Chan, P., Vasios, C.E., Bonmassar, G., Ito, N., Náñez, J.E., Sr., Shimojo, S., and Watanabe, T. (2009). Location-specific cortical activation changes during sleep after training for perceptual learning. Curr Biol 19, 1278–1282. 10.1016/j.cub.2009.06.011.

27. Bang, J.W., Shibata, K., Frank, S.M., Walsh, E.G., Greenlee, M.W., Watanabe, T., and Sasaki, Y. (2018). Consolidation and reconsolidation share behavioral and neurochemical mechanisms. Nat Hum Behav 2, 507–513. 10.1038/s41562-018-0366-8.

28. Shibata, K., Sasaki, Y., Bang, J.W., Walsh, E.G., Machizawa, M.G., Tamaki, M., Chang, L.H., and Watanabe, T. (2017). Overlearning hyperstabilizes a skill by rapidly making neurochemical processing inhibitory-dominant. Nat Neurosci 20, 470–475. 10.1038/nn.4490.

29. Yamada, T., Watanabe, T., and Sasaki, Y. (2023). Plasticity-stability dynamics during post-training processing of learning. Trends Cogn Sci. 10.1016/j.tics.2023.09.002.

30. Tamaki, M., Watanabe, T., and Sasaki, Y. (2021). Coregistration of magnetic resonance spectroscopy and polysomnography for sleep analysis in human subjects. STAR Protoc 2, 100974. 10.1016/j.xpro.2021.100974.

31. Dinges, D.F., and Powell, J.W. (1985). Microcomputer analyses of performance on a portable, simple visual RT task during sustained operations. Behavior Research Methods, Instruments, & Computers 17, 652–655. 10.3758/BF03200977.

32. Hoddes, E., Zarcone, V., Smythe, H., Phillips, R., and Dement, W.C. (1973). Quantification of sleepiness: a new approach. Psychophysiology 10, 431–436. 10.1111/j.1469-8986.1973.tb00801.x.

33. Hensch, T.K. (2005). Critical period plasticity in local cortical circuits. Nat Rev Neurosci 6, 877–888. 10.1038/nrn1787.

34. Lee, S.H., Kwan, A.C., Zhang, S., Phoumthipphavong, V., Flannery, J.G., Masmanidis, S.C., Taniguchi, H., Huang, Z.J., Zhang, F., Boyden, E.S., et al. (2012). Activation of specific interneurons improves V1 feature selectivity and visual perception. Nature 488, 379–383. 10.1038/nature11312.

35. Saper, C.B., Scammell, T.E., and Lu, J. (2005). Hypothalamic regulation of sleep and circadian rhythms. Nature 437, 1257–1263. 10.1038/nature04284.

36. Nofzinger, E.A., Buysse, D.J., Germain, A., Price, J.C., Miewald, J.M., and Kupfer, D.J. (2004). Functional neuroimaging evidence for hyperarousal in insomnia. Am J Psychiatry 161, 2126–2128. 10.1176/appi.ajp.161.11.2126.

37. Czeisler, C.A., and Gooley, J.J. (2007). Sleep and circadian rhythms in humans. Cold Spring Harb Symp Quant Biol 72, 579–597. 10.1101/sqb.2007.72.064.

38. Chee, M.W., and Chuah, L.Y. (2008). Functional neuroimaging insights into how sleep and sleep deprivation affect memory and cognition. Curr Opin Neurol 21, 417–423. 10.1097/WCO.0b013e3283052cf7.

39. Li, W., Ma, L., Yang, G., and Gan, W.B. (2017). REM sleep selectively prunes and maintains new synapses in development and learning. Nat Neurosci 20, 427–437. 10.1038/nn.4479.

40. Carskadon, M.A., and Dement, W.C. (1981). Cumulative effects of sleep restriction on daytime sleepiness. Psychophysiology 18, 107–113. 10.1111/j.1469-8986.1981.tb02921.x.

41. Abdellahi, M.E.A., Koopman, A.C.M., Treder, M.S., and Lewis, P.A. (2023). Targeting targeted memory reactivation: Characteristics of cued reactivation in sleep. Neuroimage 266, 119820. 10.1016/j.neuroimage.2022.119820.

42. Klinzing, J.G., Niethard, N., and Born, J. (2019). Mechanisms of systems memory consolidation during sleep. Nat Neurosci 22, 1598–1610. 10.1038/s41593-019-0467-3.

43. Oudiette, D., and Paller, K.A. (2013). Upgrading the sleeping brain with targeted memory reactivation. Trends Cogn Sci 17, 142–149. 10.1016/j.tics.2013.01.006.

44. Stickgold, R. (2013). Parsing the role of sleep in memory processing. Curr Opin Neurobiol 23, 847–853. 10.1016/j.conb.2013.04.002.

45. Uji, M., and Tamaki, M. (2023). Sleep, learning, and memory in human research using noninvasive neuroimaging techniques. Neurosci Res 189, 66–74. 10.1016/j.neures.2022.12.013.

46. Kuriyama, K., Stickgold, R., and Walker, M.P. (2004). Sleep-dependent learning and motor-skill complexity. Learn Mem 11, 705–713. 10.1101/lm.76304.

47. Manoach, D.S., Thakkar, K.N., Stroynowski, E., Ely, A., McKinley, S.K., Wamsley, E., Djonlagic, I., Vangel, M.G., Goff, D.C., and Stickgold, R. (2010). Reduced overnight consolidation of procedural learning in chronic medicated schizophrenia is related to specific sleep stages. J Psychiatr Res 44, 112–120. 10.1016/j.jpsychires.2009.06.011.

48. Tamaki, M., Matsuoka, T., Nittono, H., and Hori, T. (2008). Fast sleep spindle (13-15 hz) activity correlates with sleep-dependent improvement in visuomotor performance. Sleep 31, 204–211. 10.1093/sleep/31.2.204.

49. TAMAKI, M., NITTONO, H., and HORI, T. (2007). Efficacy of overnight sleep for a newly acquired visuomotor skill. Sleep and Biological Rhythms 5, 111–116. 10.1111/j.1479-8425.2007.00260.x.

50. Araj-Khodaei, M., Sanaie, S., Nejadghaderi, S.A., Sullman, M.J.M., Samei-Sis, S., Taheri-Targhi, S., Yousefi, Z., Matlabi, H., Safiri, S., and Azizi-Zeinalhajlou, A. (2022). Profile of Tabriz Older People Health Survey (TOPS-2019): a representative community-based cross-sectional study. Sci Rep 12, 17879. 10.1038/s41598-022-22710-2.

51. Mander, B.A., Winer, J.R., and Walker, M.P. (2017). Sleep and Human Aging. Neuron 94, 19–36. 10.1016/j.neuron.2017.02.004.

52. Manoach, D.S., and Stickgold, R. (2019). Abnormal Sleep Spindles, Memory Consolidation, and Schizophrenia. Annu Rev Clin Psychol 15, 451–479. 10.1146/annurev-clinpsy-050718-095754.

53. Matsui, K., Yoshiike, T., Nagao, K., Utsumi, T., Tsuru, A., Otsuki, R., Ayabe, N., Hazumi, M., Suzuki, M., Saitoh, K., et al. (2021). Association of Subjective Quality and Quantity of Sleep with Quality of Life among a General Population. Int J Environ Res Public Health 18. 10.3390/ijerph182312835.

54. Miner, B., and Kryger, M.H. (2020). Sleep in the Aging Population. Sleep Med Clin 15, 311–318. 10.1016/j.jsmc.2020.02.016.

55. Berard, A.V., Cain, M.S., Watanabe, T., and Sasaki, Y. (2015). Frequent video game players resist perceptual interference. PLoS One 10, e0120011. 10.1371/journal.pone.0120011.

56. Li, R., Polat, U., Makous, W., and Bavelier, D. (2009). Enhancing the contrast sensitivity function through action video game training. Nat Neurosci 12, 549–551. 10.1038/nn.2296.

57. Karni, A., and Sagi, D. (1991). Where practice makes perfect in texture discrimination: evidence for primary visual cortex plasticity. Proc Natl Acad Sci U S A 88, 4966–4970. 10.1073/pnas.88.11.4966.

58. Horikawa, T., Tamaki, M., Miyawaki, Y., and Kamitani, Y. (2013). Neural decoding of visual imagery during sleep. Science 340, 639–642. 10.1126/science.1234330.

59. Ehlers, C.L., and Kupfer, D.J. (1997). Slow-wave sleep: do young adult men and women age differently? J Sleep Res 6, 211–215. 10.1046/j.1365-2869.1997.00041.x.

60. Buysse, D.J., Reynolds, C.F., 3rd, Monk, T.H., Berman, S.R., and Kupfer, D.J. (1989). The Pittsburgh Sleep Quality Index: a new instrument for psychiatric practice and research. Psychiatry Res 28, 193-213. 10.1016/0165-1781(89)90047-4.

61. Horne, J.A., and Ostberg, O. (1976). A self-assessment questionnaire to determine morningness-eveningness in human circadian rhythms. Int J Chronobiol 4, 97–110.

62. Oldfield, R.C. (1971). The assessment and analysis of handedness: the Edinburgh inventory. Neuropsychologia 9, 97–113. 10.1016/0028-3932(71)90067-4.

63. Berry, R.B., Brooks, R., Gamaldo, C., Harding, S.M., Lloyd, R.M., Quan, S.F., Troester, M.T., and Vaughn, B.V. (2017). AASM Scoring Manual Updates for 2017 (Version 2.4). J Clin Sleep Med 13, 665–666. 10.5664/jcsm.6576.

64. Murphy, P.J., and Campbell, S.S. (1997). Nighttime drop in body temperature: a physiological trigger for sleep onset? Sleep 20, 505–511. 10.1093/sleep/20.7.505.

65. Basner, M., and Dinges, D.F. (2011). Maximizing sensitivity of the psychomotor vigilance test (PVT) to sleep loss. Sleep 34, 581–591. 10.1093/sleep/34.5.581.

66. Loh, S., Lamond, N., Dorrian, J., Roach, G., and Dawson, D. (2004). The validity of psychomotor vigilance tasks of less than 10-minute duration. Behav Res Methods Instrum Comput 36, 339–346. 10.3758/bf03195580.

67. Tamaki, M., Berard, A.V., Barnes-Diana, T., Siegel, J., Watanabe, T., and Sasaki, Y. (2020). Reward does not facilitate visual perceptual learning until sleep occurs. Proc Natl Acad Sci U S A 117, 959–968. 10.1073/pnas.1913079117.

68. Wang, Z., Tamaki, M., Frank, S.M., Shibata, K., Worden, M.S., Yamada, T., Kawato, M., Sasaki, Y., and Watanabe, T. (2021). Visual perceptual learning of a primitive feature in human V1/V2 as a result of unconscious processing, revealed by decoded functional MRI neurofeedback (DecNef). J Vis 21, 24. 10.1167/jov.21.8.24.

69. Yotsumoto, Y., Watanabe, T., and Sasaki, Y. (2008). Different dynamics of performance and brain activation in the time course of perceptual learning. Neuron 57, 827–833. 10.1016/j.neuron.2008.02.034.

70. Yotsumoto, Y., Chang, L.H., Watanabe, T., and Sasaki, Y. (2009). Interference and feature specificity in visual perceptual learning. Vision Res 49, 2611–2623. 10.1016/j.visres.2009.08.001.

71. Karni, A., and Sagi, D. (1993). The time course of learning a visual skill. Nature 365, 250–252. 10.1038/365250a0.

72. Machizawa, M., Patey, R., Kim, D., and Watanabe, T. (2014). Different aspects of training on a texture discrimination task (TDT) improves different attentional abilities. Journal of Vision 14, 951–951.

73. Brainard, D.H. (1997). The Psychophysics Toolbox. Spat Vis 10, 433–436.

74. Pelli, D.G. (1997). The VideoToolbox software for visual psychophysics: transforming numbers into movies. Spat Vis 10, 437–442.

75. Wichmann, F.A., and Hill, N.J. (2001). The psychometric function: I. Fitting, sampling, and goodness of fit. Percept Psychophys 63, 1293–1313. 10.3758/bf03194544.

76. Edden, R.A., and Barker, P.B. (2007). Spatial effects in the detection of gamma-aminobutyric acid: improved sensitivity at high fields using inner volume saturation. Magn Reson Med 58, 1276–1282. 10.1002/mrm.21383.

77. Mescher, M., Merkle, H., Kirsch, J., Garwood, M., and Gruetter, R. (1998). Simultaneous in vivo spectral editing and water suppression. NMR Biomed 11, 266–272. 10.1002/(sici)1099-1492(199810)11:6<266::aid-nbm530>3.0.co;2-j.

78. Provencher, S.W. (1993). Estimation of metabolite concentrations from localized in vivo proton NMR spectra. Magn Reson Med 30, 672–679. 10.1002/mrm.1910300604.

79. Provencher, S.W. (2001). Automatic quantitation of localized in vivo 1H spectra with LCModel. NMR Biomed 14, 260–264. 10.1002/nbm.698.

